# Carbapenem- and cefiderocol-resistant *Enterobacterales* in surface waters in Kumasi, Ashanti Region, Ghana

**DOI:** 10.1101/2023.08.04.551947

**Authors:** Elias Eger, Timo Homeier-Bachmann, Eugene Adade, Sylvia Dreyer, Stefan E. Heiden, Patrick Ofori Tawiah, Augustina Angelina Sylverken, Sascha Knauf, Katharina Schaufler

## Abstract

This study investigated the presence of extended-spectrum β-lactamase-producing *Enterobacterales* in selected surface waters in the Ashanti Region in Ghana. Subsequent genomic analysis revealed that one-fifth of the isolates were genotypically carbapenem-resistant. Phenotypic susceptibility testing not only confirmed their carbapenem resistance, but also uncovered two cefiderocol-resistant isolates.

## MAIN

Antimicrobial resistance (AMR) has emerged as an urgent global health crisis, threatening the effectiveness of life-saving antibiotics, including carbapenems (e.g., ertapenem and meropenem). Sub-Saharan Africa (SSA) bears the greatest burden of AMR, with a significant number of deaths attributed to carbapenem-resistant *Enterobacterales* (CR-E) (e.g., *Escherichia coli, Enterobacter* spp., and *Klebsiella* spp.) [1]. The global spread of CR-E, traditionally considered nosocomial pathogens, demonstrates their ability to cross species boundaries and spread at the human-animal-environmental interface. Ghana, like many other low- and middle-income countries faces challenges with clean water, limited sanitation and health care, a lack of public education and environmental consciousness. Together, these factors facilitate the spread and persistence of CR-E within human clinics and in animal and environmental reservoirs, including surface water. However, the epidemiology of CR-E in Ghanaian surface water remains largely unexplored.

This study aimed to fill this gap by investigating the presence of CR-E in surface waters from Kumasi, the capital city of the Ashanti Region of Ghana. Kumasi is characterized by a diverse population and various peri-urban water sources used by humans and animals. It therefore provides an ideal setting to assess the distribution of CR-E in surface water in a typical West African cityscape, to analyze AMR genetics, and to investigate the susceptibility profiles of CR-E isolates.

### The Study

The study’s objective was to complement the Surveillance Outbreak Response Management and Analysis System (SORMAS; [2]) with data on the occurrence of antimicrobial-resistant pathogens. In this frame, we collected water samples from ten peri-urban sites in the Kumasi area weekly from week 31 (July) to week 38 (September) 2021 (Figure 1). Overall, we isolated 121 extended-spectrum β-lactamase (ESBL)-producing *Enterobacterales* and *Pseudomonas aeruginosa*. The most prevalent species was *E. coli*, accounting for 70.25% (85/121) of the isolates, followed by *K. pneumoniae* sensu stricto (hereafter referred to as *K. pneumoniae*; 23.14%; 28/121), *E. cloacae* (4.13%; 5/121), and *P. aeruginosa* (0.83%; 1/121). All isolates were whole-genome sequenced on an Illumina NextSeq 2000 platform, followed by bioinformatics downstream analysis based on previously published protocols [3]. AMR gene analysis using the AMRFinderPlus database [4] showed that all isolates carried ESBL genes, with *bla*_CTX-M-15_ as the dominant gene (94.21%; 114/121). Alarmingly, 23 (19.01%; 23/121) isolates carried genes associated with carbapenem resistance. Specifically, 15 *E. coli*, six *K. pneumoniae*, and two *E. cloacae* were positive for carbapenemase-encoding genes, with the predominant gene being *bla*_OXA-181_ (69.57%; 16/23), followed by *bla*_NDM-5_ (21.74%; 5/23), and *bla*_OXA-48_ (8.70%; 2/23).

**Figure 1:**
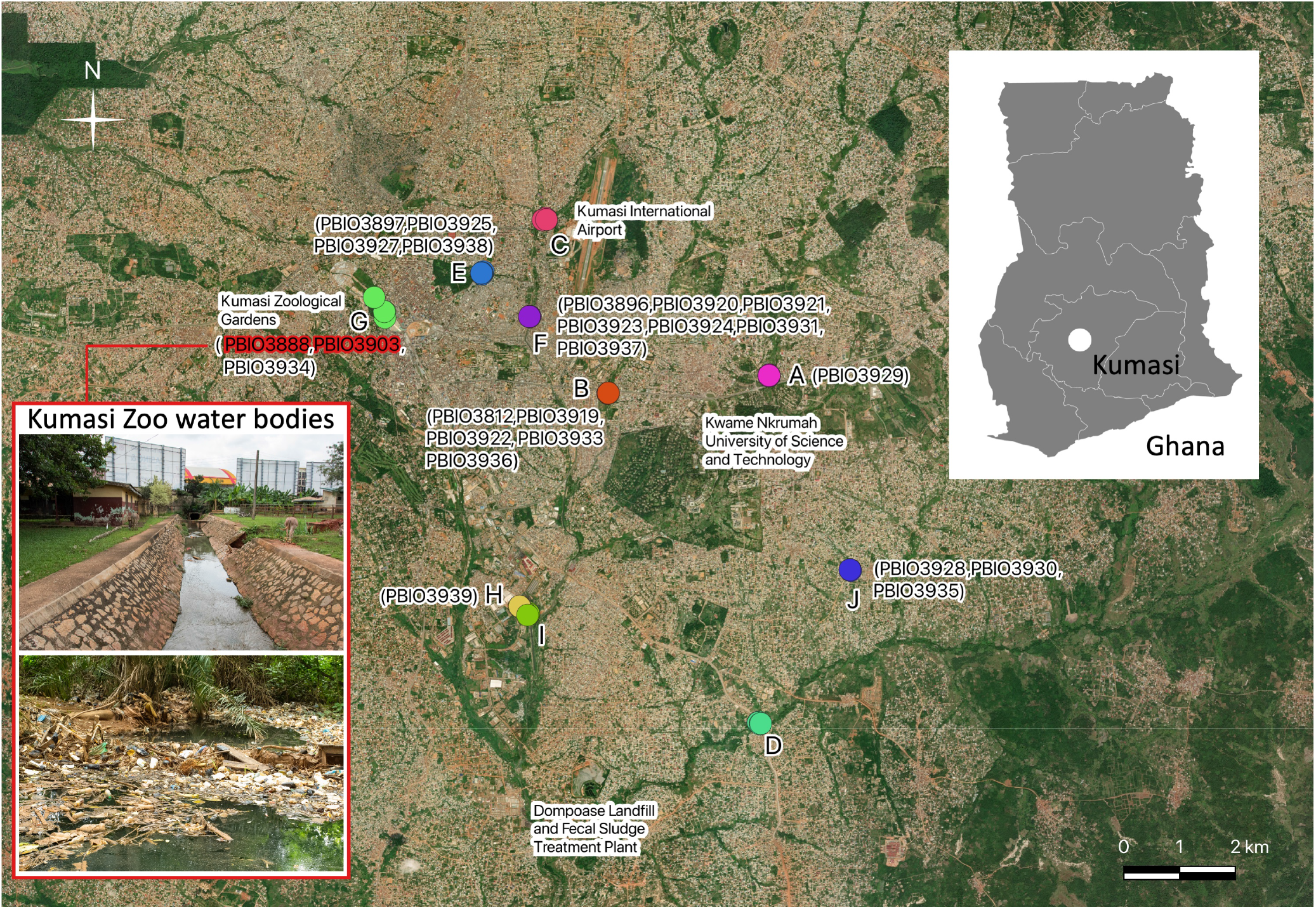
Map of the Kumasi area (Ghana). The selected surface waters are marked in different colors and numbered from A to J. The CR-E isolates are assigned to their different sampling locations. The inset image outlined in red shows images of the Kumasi Zoo water bodies (location G) where the two cefiderocol-resistant isolates (highlighted in red) were isolated. For reference, the overall location of Kumasi is indicated on the Ghana overview map and included as a smaller inset.

Given the clinical significance of CR-E, we performed phenotypic antimicrobial susceptibility testing for these isolates using the automated VITEK 2 system (AST-N428 and AST-XN24; bioMérieux, Marcy l’Etoile, France) to assess carbapenem-resistant phenotypes. In addition, we evaluated susceptibility to the recently approved and important last-resort siderophore cephalosporin cefiderocol using disk diffusion assays with cefiderocol 30 μg disks (Mast Diagnostics, Merseyside, UK). Isolates falling within the area of technical uncertainty (*Enterobacterales*, 18–22 mm [5]) were retested using a commercial broth microdilution kit (ComASP, Liofilchem, Waltham, Massachusetts, USA) according to the manufacturer’s instructions. We also included the *P. aeruginosa* isolate PBIO3812, as clinical *P. aeruginosa* isolates exhibit reduced membrane permeability, leading to reduced sensitivity to carbapenems [6]. All results were interpreted according to the published breakpoints of EUCAST [5] and are summarized in Table 1.

**Table 1:** Overview of the CR-E and their phenotypic and genotypic properties.

| Isolate | Species | ST | Carbapenemase<br>gene<sup>a</sup> | Phenotype |  |  |  |  |  |  |  |  |
| --- | --- | --- | --- | --- | --- | --- | --- | --- | --- | --- | --- | --- |
|  |  |  |  | ETP |  | IPM |  | MEM |  | ZD<br>[mm] | FDC<br>MIC<br>[μg/mL] | S/R<sup>b</sup> |
|  |  |  |  | MIC<br>[μg/mL] | S/I/R<sup>b</sup> | MIC<br>[μg/mL] | S/I/R<sup>b</sup> | MIC<br>[μg/mL] | S/I/R<sup>b</sup> |  |  |  |
| PBIO3812 | PAe | 463 | none | ND | N/A | 2 | I | ≤0.25 | S | 31 | ND | S |
| PBIO3888 | ECo | 1588 | *bla*<sub>NDM-5</sub> | ≥8 | R | ≥16 | R | ≥16 | R | 19<sup>c</sup> | 8<sup>c</sup> | R |
| PBIO3896 | ECo | 410 | *bla*<sub>NDM-5</sub> | ≥8 | R | ≥16 | R | ≥16 | R | 20<sup>c</sup> | 2<sup>c</sup> | S |
| PBIO3897 | ECo | 44 | *bla*<sub>OXA-181</sub> | 1 | R | 1 | S | 0.5 | S | 24 | ND | S |
| PBIO3903 | ECo | 1588 | *bla*<sub>NDM-5</sub> | ≥8 | R | ≥16 | R | ≥16 | R | 19<sup>c</sup> | 4<sup>c</sup> | R |
| PBIO3919 | KPn | 2668 | *bla*<sub>OXA-181</sub> | 2 | R | 4 | I | 2 | S | 28 | ND | S |
| PBIO3920 | KPn | 25 | *bla*<sub>OXA-181</sub> | 2 | R | 4 | I | 2 | S | 28 | ND | S |
| PBIO3921 | ECl | 171 | *bla*<sub>OXA-48</sub> | 2 | R | 4 | I | 4 | I | 24 | ND | S |
| PBIO3922 | KPn | 2668 | *bla*<sub>OXA-181</sub> | ≥8 | R | 4 | I | 2 | S | 29 | ND | S |
| PBIO3923 | KPn | 25 | *bla*<sub>OXA-181</sub> | 2 | R | 4 | I | 2 | S | 27 | ND | S |
| PBIO3924 | ECl | 171 | *bla*<sub>OXA-48</sub> | 1 | R | 4 | I | 4 | I | 24 | ND | S |
| PBIO3925 | KPn | 25 | *bla*<sub>OXA-181</sub> | 1 | R | 2 | S | 2 | S | 28 | ND | S |
| PBIO3927 | KPn | 25 | *bla*<sub>OXA-181</sub> | 2 | R | 2 | S | 2 | S | 28 | ND | S |
| PBIO3928 | ECo | 410 | *bla*<sub>OXA-181</sub> | 4 | R | 0.5 | S | 0.5 | S | 22 | ND | S |
| PBIO3929 | ECo | 410 | *bla*<sub>OXA-181</sub> | 2 | R | 0.5 | S | 2 | S | 23 | ND | S |
| PBIO3930 | ECo | 6359 | *bla*<sub>OXA-181</sub> | 1 | R | 1 | S | ≤0.25 | S | 25 | ND | S |
| PBIO3931 | ECo | 410 | *bla*<sub>OXA-181</sub> | 2 | R | 0.5 | S | 2 | S | 22 | ND | S |
| PBIO3933 | ECo | 410 | *bla*<sub>OXA-181</sub> | 4 | R | 1 | S | 2 | S | 23 | ND | S |
| PBIO3934 | ECo | 1588 | *bla*<sub>NDM-5</sub> | ≥8 | R | ≥16 | R | ≥16 | R | 20<sup>c</sup> | 2<sup>c</sup> | S |
| PBIO3935 | ECo | 410 | *bla*<sub>OXA-181</sub> | 4 | R | 0.5 | S | 2 | S | 23 | ND | S |
| PBIO3936 | ECo | 410 | *bla*<sub>NDM-5</sub> | ≥8 | R | 8 | R | ≥16 | R | 26 | ND | S |
| PBIO3937 | ECo | 6359 | *bla*<sub>OXA-181</sub> | 4 | R | 0.5 | S | 2 | S | 23 | ND | S |
| PBIO3938 | ECo | 6359 | *bla*<sub>OXA-181</sub> | 2 | R | 1 | S | 2 | S | 29 | ND | S |
| PBIO3939 | ECo | 410 | *bla*<sub>OXA-181</sub> | ≥8 | R | 1 | S | 2 | S | 22 | ND | S |

<sup>a</sup>Predictions for carbapenemase genes are based on alignments of sequences from the AMRFinderPlus database [4] (default settings of identity [use a curated threshold if it exists, and ≥90.0% otherwise] and coverage [≥50.0%]). <sup>b</sup>Interpretation categories according to EUCAST guidelines [5]. <sup>c</sup>For isolates with zone diameters within the range of technical uncertainty (*Enterobacterales*, 18–22mm [5]), the minimum inhibitory concentration of cefiderocol was determined by broth microdilution. ECl, *E. cloacae*; ECo, *E. coli*; ETP, ertapenem; FDC, cefiderocol; I, susceptible, increased exposure; IPM, imipenem; KPn, *K. pneumoniae*; MEM, meropenem; MIC, minimum inhibitory concentration; N/A, not applicable; ND, not determined; PAe, *P. aeruginosa* R, resistant; S, susceptible, standard dosing regimen; ST, sequence type; ZD, zone diameter.

## Conclusions

The *bla*_OXA-181_-positive isolates were only resistant to ertapenem, highlighting the weak hydrolytic activity of this OXA-48-like carbapenemase against carbapenems [7]. In contrast, isolates carrying *bla*_NDM-5_ or *bla*_OXA-48_ showed higher MIC values for imipenem and/or meropenem. Notably, the CR-E were predominantly associated with internationally recognized high-risk clonal lineages, such as *E. cloacae* sequence type (ST)171, *E. coli* ST410 and ST1588, and *K. pneumoniae* ST25. Studies conducted by us and others have consistently demonstrated that these clonal lineages harbor multiple AMR determinants, can be rapidly transmitted among and persist in different host species and ecosystems, are capable of causing severe disease in animals and humans, and are globally distributed (e.g., [8]). Evidence suggests the epidemiologic success of such high-risk clonal lineages results from a sophisticated interplay of factors, such as high bacterial fitness and virulence and transmission dynamics [3, 8, 9]. However, despite the growing challenges posed by AMR in SSA, this region has not been studied as extensively as others. Hence, the findings of this study significantly contribute to the understanding of CR-E epidemiology in selected surface waters in Kumasi and provide valuable insights for integrated surveillance systems such as SORMAS.

Like many other countries in SSA, Ghana faces challenge of environmental contamination with heavy metals, particularly those associated with illegal gold mining activities [10]. The presence of heavy metals in the environment harms bacteria by disrupting their physiological processes and has the potential for co-selection of AMR. This co-selection results from coupling resistance mechanisms to both antibiotics and heavy metals, with different resistance determinants often found on the same genetic element [11]. Except for seven of the *E. coli* isolates in this study, all CR-E were positive for genes associated with multimetal RND efflux pump activity (*silABCEFPRS*), which typically confer resistance to various heavy metals, including copper and silver. The predominant genotype among the heavy metal-resistant isolates was associated with copper resistance (*pcoABCDRS*; 17/24), followed by arsenic resistance (*arsD*; 8/24), mercury resistance (*merRT*; 7/24), and tellurium resistance (*terD*; 4/24). The co-existence of heavy metal resistance and carbapenemase genes in these bacteria underscores the potential for co-selection and complicates efforts to combat both environmental contamination and the spread of AMR in this region in SSA. Consequently, assessing heavy metal resistance determinants should be an elementary component of surface water surveillance approaches [12]. Future studies in Kumasi should include measures of heavy metal contamination of surface waters to follow up on the possible contributing drivers of AMR dissemination.

To our knowledge, this is the first report describing cefiderocol-resistant *Enterobacterales* in surface waters from SSA. Notably, both resistant isolates, PBIO3888 and PBIO3903, were obtained shortly after the international clinical use of cefiderocol. Recent studies have proposed several mechanisms contributing to cefiderocol resistance, including gene alterations in the iron transport pathway and nutrient uptake (e.g., *cirA* and *ompC*) [13]. However, a BLAST analysis of the amino acid sequences of CirA (UniProt accession P17315), OmpF (UniProt accession P02931), and OmpC (UniProt accession P06996), using *E. coli* K-12 as a reference, did not reveal any potential resistance-mediating mutations. This suggests that cefiderocol resistance may be attributed to overexpression of the NDM-5 carbapenemase. A recent study by Nurjadi *et al*. showed that the carriage of *bla*_NDM-5_ is a “genetic risk factor” that facilitates the development of resistance to cefiderocol [14]. However, further research is needed to identify the underlying mechanisms of cefiderocol resistance in our samples. Nevertheless, the occurrence of cefiderocol-resistant *Enterobacterales* in surface waters (i.e., in the environment) of Kumasi in Ghana is alarming.

In conclusion, understanding the resistance epidemiology of CR-E in different environments is crucial for effective surveillance and management of AMR in SSA. Integrating genomic and epidemiologic data, combined with cutting-edge surveillance solutions (e.g., SORMAS), will facilitate early identification, monitoring of resistance patterns, identication of high-risk areas, development of effective prevention strategies, and information of health authorities. Also, this study emphasizes the importance of a multidisciplinary One Health approach that considers human, animal, and environmental health by addressing the complex challenges AMR poses in SSA.

## NOTES

### Author contributions

Conceptualization, K.S. and S.K.; methodology, A.S., E.A., P.O.T. and S.D.; software, E.E. and S.E.H.; validation, E.E. and T.H.-B; formal analysis, A.S., E.E., K.S., S.E.H., and S.K.; investigation, A.S., E.A., E.E., S.D., P.O.T. and T.H.-B; resources, A.S., K.S., S.K. and T.H.-B; data curation, S.E.H.; writing—original draft preparation, E.E. and K.S.; writing—review and editing, A.S., E.A., E.E., K.S., P.O.T., S.D., S.K. and T.H.-B.; visualization, E.E., S.K. and S.E.H.; supervision, K.S. and S.K.; project administration, S.K.; funding acquisition, A.S., K.S., S.K. and T.H.-B. All authors have read and agreed to the published version of the manuscript.

## Acknowledgments

We thank Sara-Lucia Wawrzyniak (Pharmaceutical Microbiology, Institute of Pharmacy, University of Greifswald) and Simone Lueert (Institute of International Animal Health/One Health, Friedrich-Loeffler-Institut) for their excellent technical assistance.

## Data availability

The data for this study have been deposited in the European Nucleotide Archive (ENA) at EMBL-EBI under accession number PRJEB64632 (https://www.ebi.ac.uk/ena/browser/view/PRJEB64632).

## Funding

This work was supported by a seed grant from the Helmholtz Centre for Infection Research (HZI, Germany) entitled “Water-based outbreak prediction in peri-urban Africa”. Support was also obtained from a grant from the Federal Ministry of Education and Research (BMBF, Germany) to KS entitled “Disarming pathogens as a different strategy to fight antimicrobial-resistant Gram-negatives” (01KI2015).

## Potential conflicts of interest

All authors report no potential conflicts.

## Notes

### Competing Interest Statement

The authors have declared no competing interest.

